# Time-resolved transcriptomic profiling of *Dictyostelium discoideum* infection with *Mycobacterium marinum* reveals Atg9-dependent restriction via maintenance of vacuole integrity

**DOI:** 10.64898/2026.09.10.750649

**Authors:** Jahn Nitschke, Justine Toinon, Frédéric Burdet, Astrid Melotti, Hubert Hilbi, Pierre Cosson, Marco Pagni, Thierry Soldati, Nabil Hanna

**Author notes:** Address correspondence to Thierry Soldati, and Nabil Hanna.

## Abstract

Tuberculosis (TB), caused by *Mycobacterium tuberculosis* (Mtb), remains a major global health challenge, highlighting the need to better understand host mechanisms restricting mycobacteria infection. *Mycobacterium marinum* (Mm) shares key virulence mechanisms with Mtb and provides a suitable model for studying mycobacterial pathogenesis. Here, we used *Dictyostelium discoideum* (Dd), a genetically tractable phagocytic model with evolutionarily conserved pathways shared with mammalian macrophages, to investigate host responses to Mm infection. Time-resolved transcriptomic profiling across early, intermediate, and late stages of infection identified global infection-responsive genes, stage-specific pathways, and substantial conservation with transcriptional responses of human macrophages infected with Mtb. Autophagy and ESCRT pathways were dynamically regulated throughout infection, suggesting stage-specific roles in host defence. We focused on Atg9, an autophagy factor that was strongly induced during infection and recruited to damaged mycobacterium-containing vacuoles (MCVs). Atg9 promoted membrane damage control and maintained MCV integrity, thereby preventing premature bacteria escape to the cytosol of virulent Mm. Loss of Atg9 led to accumulation of MCV damage, accelerated escape to the cytosol, and enhanced intracellular bacteria growth, phenocopying *atg*1-deficient cells. Our findings further distinguish complementary host defence mechanisms acting at distinct stages of infection: Atg9-associated membrane repair, potentially involving ATG8ylation, limits vacuole damage and bacteria escape during early infection, whereas Atg9-dependent xenophagic restriction contributes to bacteria clearance at later stages. Together, these findings establish Dd as a powerful model for dissecting conserved host responses to mycobacteria and identify Atg9-dependent membrane protection as a key host resistance mechanism with potential relevance to TB pathogenesis and therapeutic intervention.

**IMPORTANCE:** *M. tuberculosis* (Mtb) infection remains a major global health concern. Using *Dictyostelium discoideum* (Dd) as a model to study *Mycobacterium marinum* (Mm) infection, this study identifies evolutionarily conserved host pathways relevant to mammalian macrophages. Time-resolved transcriptomic profiling reveals both global and stage-specific responses to infection, including dynamic regulation of the autophagy and ESCRT machineries, which play central roles in host defence against intracellular pathogens. Importantly, we identify Atg9 as a host resistance factor that maintains the integrity of mycobacterium-containing vacuoles (MCVs) and prevents premature bacteria escape to the cytosol. More broadly, our findings reveal that membrane repair and xenophagic bacteria restriction constitute complementary layers of host defence that act at distinct stages of infection. The conservation of infection-induced transcriptional responses between Dd and human macrophages further supports the value of this model for identifying host pathways relevant to tuberculosis and provides a framework for exploring host-directed therapeutic strategies.

## INTRODUCTION

Tuberculosis (TB), caused by *Mycobacterium tuberculosis* (Mtb), remains a major global health threat. *Mycobacterium marinum* (Mm), is an environmental, opportunistic pathogen of ectotherms and humans, shares key virulence mechanisms with Mtb and is an established model for studying mycobacterial pathogenesis (1–3). The social amoeba *Dictyostelium discoideum* (Dd) provides a genetically tractable model of professional phagocytes, with conserved phagosomal and cell-autonomous defence pathways shared with mammalian macrophages (3–5). Together, the Dd–Mm system enables mechanistic investigation of host– mycobacterium interactions using genetic, biochemical, and live-cell imaging approaches.

Mm infection of Dd progresses through distinct stages, beginning with manipulation of phagosome maturation and establishment of the mycobacterium-containing vacuole (MCV) early during infection, followed by progressive membrane damage, intracellular proliferation and ultimately escape to the cytosol, continued bacteria proliferation, and finally egress (3, 6). Throughout this process, the MCV represents a dynamic host–pathogen interface in which bacterial virulence factors continuously challenge host membrane integrity (7, 8). In response, Dd deploys conserved and temporally coordinated cell-autonomous defence mechanisms, including autophagy, metabolic reprogramming, and ion homeostasis, which collectively limit bacteria replication, support membrane repair, and promote pathogen clearance (4, 9–12).

A key layer of host control operates directly at the MCV through conserved membrane repair pathways. Small-scale membrane damage at the MCV of Mtb-infected macrophages (13) and Mm-infected Dd (14) is efficiently repaired by the ESCRT machinery, thereby preserving vacuolar integrity and ensuring bacteria containment (15). In Dd, ESCRT components are actively recruited to damaged MCVs, where they assemble into discrete patches and ring-like structures at sites of membrane damage (16). Conversely, Mtb can counteract this repair response through the ESX-3 effectors EsxG and EsxH, which interfere with ESCRT-III recruitment to damaged membranes and thereby promote bacteria escape (17). Despite these advances, how membrane repair and other cell-autonomous defence pathways are coordinated throughout infection, and how their relative contributions change across infection stages, remains poorly understood.

Autophagy is a conserved degradation pathway that delivers damaged cellular components and intracellular pathogens to lysosomes (18). In host defence, canonical autophagy promotes xenophagic clearance of bacteria, while emerging evidence indicates that Atg8ylation also contributes to membrane remodelling, repair, and maintenance of compartment integrity during infection (19–21). Atg9 is the only transmembrane core autophagy protein and traffics membranes between endosomal compartments and phagophore assembly sites to support autophagosome biogenesis (22–24). Beyond its canonical role in autophagy, Atg9 has emerged as a broader regulator of host defence and membrane homeostasis. Atg9B promotes Group A *Streptococcus* internalisation through actin remodelling (25), while Atg9 restricts *Rickettsia* independently of autophagy by limiting receptor binding (26). Atg9A also promotes antibacteria autophagy of *Salmonella*-containing vacuoles (27) and trafficking of *C. neoformans*-containing compartments to degradative organelles (28). More recently, Atg9 has been recruited to damaged lysosomes in a Ca²⁺-dependent and Atg8-independent manner, implicating it directly in membrane repair (29). Together, these findings position Atg9 as a versatile regulator of cytoskeletal, autophagic, and membrane-repair responses to microbial and sterile damage.

Recent transcriptomic studies have advanced our understanding of host responses to mycobacteria infection, revealing that macrophage ontogeny and its associated transcriptional programs critically shape control of intracellular mycobacteria growth (30). In Dd, early transcriptional responses to Mm infection have been characterized and shown to share features with those of human macrophages (31). Building on these findings, we comprehensively characterized the temporal transcriptional response of Dd to Mm infection across early, intermediate, and late stages of infection, and compared these dynamics with multiple transcriptomic datasets from Mtb-infected mammalian macrophages. In parallel, we investigated the role of Atg9 in host defence, demonstrating its recruitment to MCV in a damage-dependent manner. Functional analyses further revealed that Atg9 loss results in premature bacteria access to the host cytosol and enhanced intracellular bacteria proliferation.

## RESULTS

We performed a transcriptomic time-course analysis of Dd infected with Mm (Dd–Mm) over 48h. Cells were infected with GFP-expressing Mm by spinoculation or mock-infected, then collected at 1, 3, 6, 12, 24, 36, and 48 hpi. Two complementary experiments captured infection dynamics: a high-resolution early/intermediate time course (1, 3, 6, 12 hpi) and a late time course (24, 36, 48 hpi), with 1 hpi included in both for cross-comparison. Prior to RNA extraction, infected cultures were sorted by FACS into infected (GFP⁺) and non-infected (GFP⁻) populations; GFP⁻ samples likely include both bystander cells and cells that had cleared or released bacteria prior to sorting, as previously described (14). Mock-infected cells were processed in parallel. RNA was isolated from GFP⁺, GFP⁻, and mock-infected conditions for library preparation and sequencing, with each condition/time point generated in biological triplicate. After sequencing and quality control, the early time course yielded 74 samples (1 hpi: GFP⁺ n=6, GFP⁻ n=7, mock-infected n=7; 3 hpi: n=5/5/6; 6 hpi: n=8/3/5; 12 hpi: n=8/6/8), and the late time course yielded 87 samples (1 hpi: n=8/7/3; 24 hpi: n=9/9/8; 36 hpi: n=8/6/7; 48 hpi: n=8/6/8), totalling 161 samples included in downstream analyses.

### Infection status and temporal progression are major drivers of transcriptomic variance

After low-count filtering and variance-stabilizing normalization, PCA was used to assess transcriptomic variation across samples (Fig. 1). In the early dataset (Fig. 1A–E), samples separated primarily by infection status along PC2 (Fig. 1A), independent of time; separation was more pronounced between GFP⁺ and mock-infected samples than between GFP⁻ and mock-infected, suggesting GFP⁻ cells occupy an intermediate transcriptional state. Temporal variation was captured by PC1 (Fig. 1B), with the strongest separation between 1 hpi and later time points, likely reflecting the acute response to initial pathogen encounter. Each condition analysed independently showed clear temporal progression along PC1 (Fig. 1C–E).

**Fig. 1.**
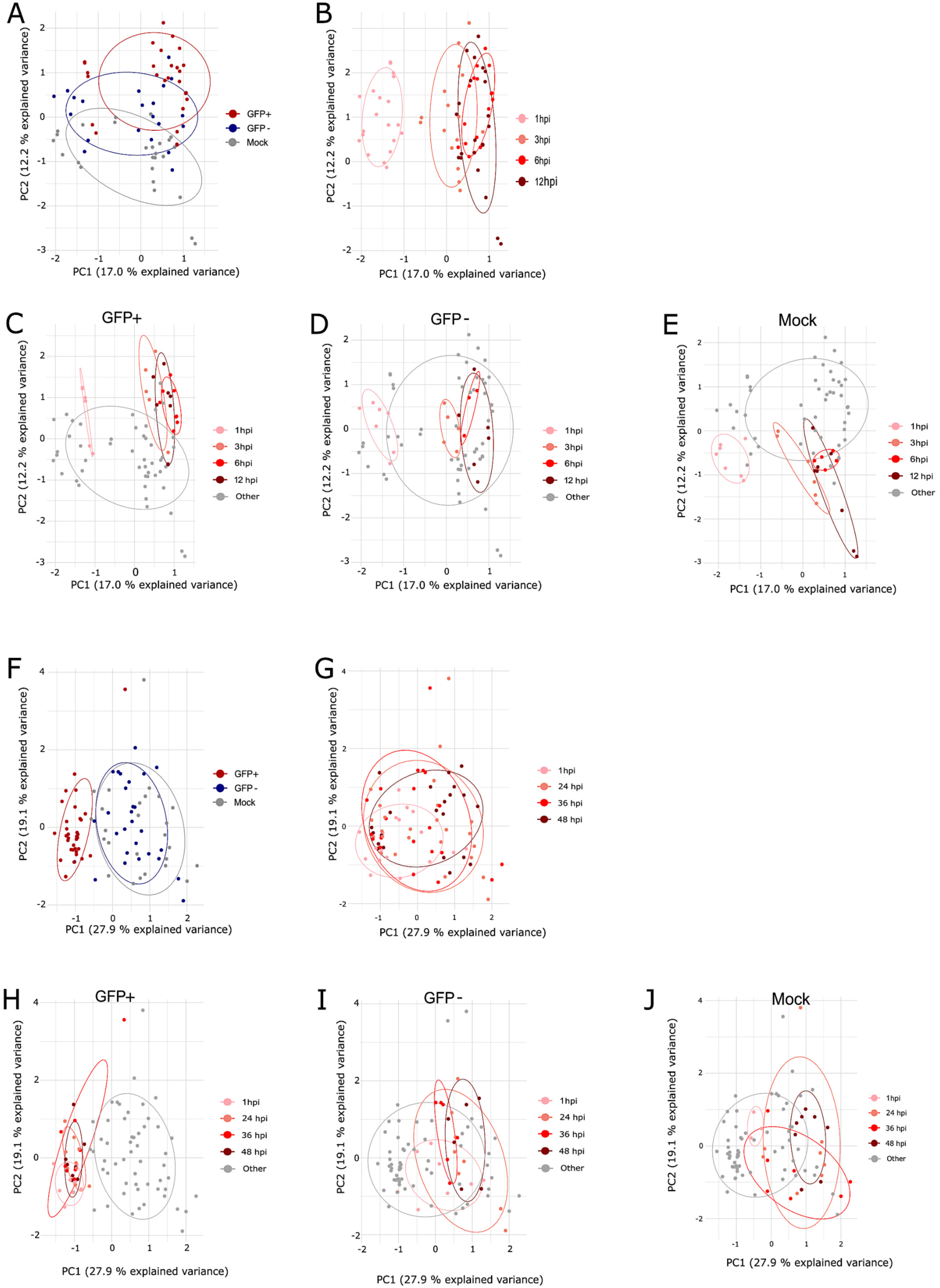
Principal Component Analysis. Principal component analysis of datasets comprising early and late samples. The early samples (1, 3, 6 and 12 hpi) are depicted in panels **A**, **B**, **C**, **D** and **E**, coloured by infection status (**A**), time point (**B**), GFP+ and time point (**C**), GFP- and time point (**D**) and Mock-infected and time point (**E**). Variance associated with time point was captured in principal component 1 (17.0 % of total variance), variance associated with infection status was captured in principal component 2 (12.2 % of total variance). The late samples (1, 24, 36, 48 hpi) are depicted in panels **F**, **G**, **H**, **I** and **J**, coloured by infection status (**F**), time point (**G**), GFP+ and time point (**H**), GFP- and time point (**I**) and Mock-infected and time point (**J**). Variance associated with infection status was captured in principal component 1 (27.9 % of total variance), variance associated with time point was mostly captured in principal component 2 (19.1 % of total variance).

PCA of the late dataset (Fig. 1F–J) also showed clustering by infection status, but here captured primarily by PC1 (Fig. 1G), indicating infection status became the dominant source of variance at later stages. GFP⁺ samples clustered distinctly from GFP⁻ and mock-infected cells, while GFP⁻ cells increasingly overlapped with mock-infected controls, suggesting reversion toward a non-infected state by 24 hpi. Temporal separation, captured along PC2, was less pronounced than in the early dataset; condition-specific PCA showed temporal variation in GFP⁺ samples mapped to PC2, while GFP⁻ and mock-infected variation mapped to PC1 (Fig. 1H–J). Although 1 hpi was included in both series for cross-comparison, high variability at this time point precluded robust alignment between datasets. Together, these analyses identify infection status and infection stage as major determinants of transcriptomic variance. Because separation was clearest and most consistent between GFP⁺ and mock-infected samples, this comparison was selected for subsequent differential expression analyses.

### Differential gene expression analysis reveals dynamic and stage-specific host responses

To define infection-induced transcriptional responses, we performed differential gene expression analysis (DEGA) comparing GFP⁺ and mock-infected cells at each time point (FDR ≤ 0.05, |Log_2_FC| ≥ 0.585) (Table S1, S2). Across both datasets, responses were dominated by gene induction rather than repression, with the largest fold changes at 1 and 6 hpi in the early dataset and at 1 hpi in the late dataset (Fig. 2A–H). Among the most strongly induced transcripts were *iliL*, *rigA*, and the poorly characterized DDB_G0281135, predicted to encode a secreted or membrane-associated protein. Conversely, udu family genes (*uduA1–A3*, *uduB*, *uduG*) previously linked to dupA-regulated MAP kinase signaling during *Legionella pneumophila* (Lp) infection were consistently downregulated (32), suggesting convergent responses to distinct intracellular pathogens.

**Fig. 2.**
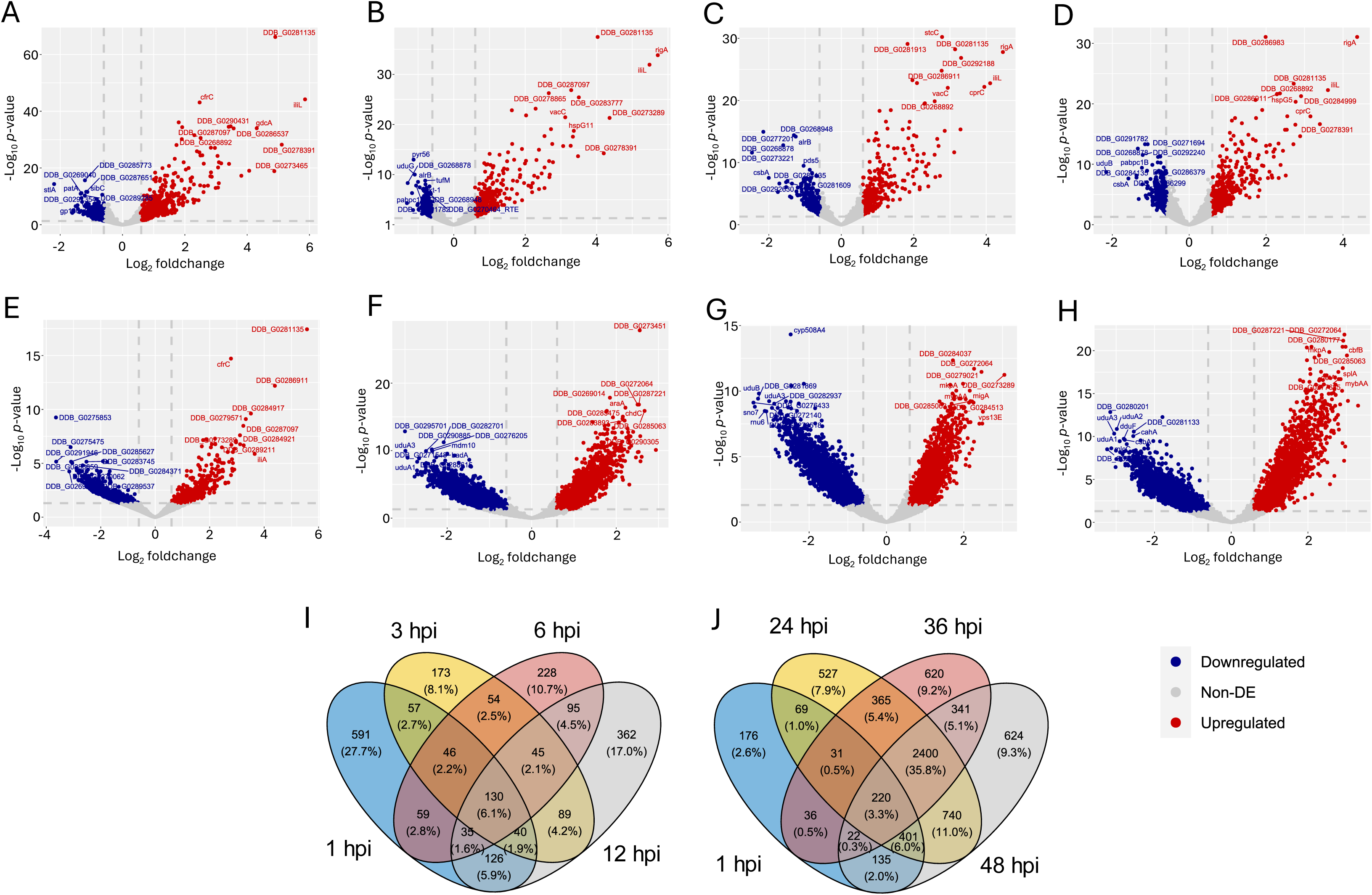
Differential Expression Analysis. Volcano plots and time point intersections of differentially expressed genes. Volcano plots of contrasts between GFP+ and Mock-infected from the early samples (1, 3, 6 and 12 hpi) are depicted in panels **A** (1 hpi), **B** (3 hpi), **C** (6 hpi), and **D** (12 hpi). Volcano plots of contrasts between GFP+ and Mock-infected from the late samples (1, 24, 36 and 48 hpi) are depicted in panels **E** (1 hpi), **F** (24 hpi), **G** (36 hpi), and **H** (48 hpi). All genes with an absolute log2 fold change ≥ 0.585 and a false discovery rate ≤ 0.05 were considered differentially expressed. The dashed lines in the volcano plots represent these thresholds. Upregulated genes are coloured in red, downregulated genes in blue, non-differentially expressed genes are coloured in grey. Additionally, the ten most pertinent up- or downregulated genes are labelled with their gene symbol or gene identifier. Panels **I** and **J** show the set analysis of both datasets. The number of differentially expressed genes common to all analysed time points within each set are depicted in the middle intersection, the number of differentially expressed genes unique to each time point are depicted at the fringes of the Venn diagrams.

Comparing DEG overlaps across time points (Fig. 2I, Table S1,S2) identified 130 genes (6.1%) consistently regulated throughout early infection and 230 genes (3.3%) throughout late infection, with 21 shared genes defining a core global signature. Early responses were dominated by rapid, time-point-specific changes (591 unique DEGs, 27.7%, at 1 hpi), whereas late infection showed a sustained program (2,400 genes, 35.8%, shared across 24–48 hpi). Ranking genes by effect size and significance, hierarchical clustering of top-ranked global and time point-specific DEGs (Fig. 3) identified distinct transcriptional signatures. The early global signature featured small heat shock proteins (*hspG8/G9/G11*), consistent with stress responses to Mm infection (Fig. 3A) (33, 34). The late global signature showed strong induction of vta1, an ESCRT component implicated in membrane repair (Fig. 3B). At 12 hpi, proteasome-associated genes (*psmD3, psmA4, psmC4, psmD13, psmB6, psmA5*) were induced, particularly in GFP⁺ cells, alongside transient atg12 induction and sustained *cdk1* downregulation, indicating early autophagy activation and cell-cycle modulation (Fig. 3C). Among late-specific markers, *zntB* was strongly induced, consistent with zinc-mediated restriction of Mm, while sustained *vacC* induction contrasted with transient *vacA*/*vacB* regulation, suggesting dynamic vacuolin-family membrane remodelling (Fig. 3D) (11, 12, 15, 35).

**Fig. 3.**
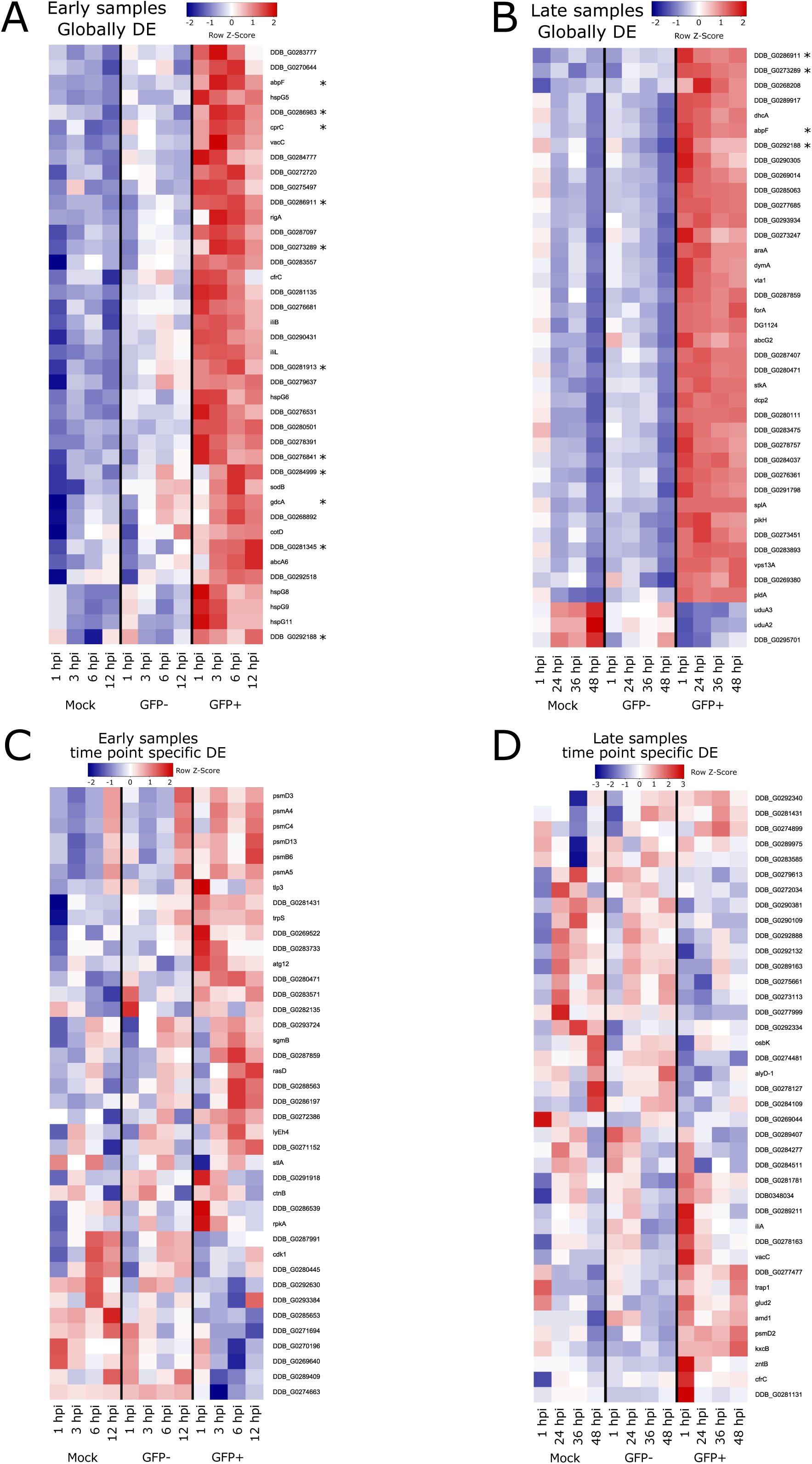
Global and time point specific markers. Time evolution of most dominant genes in the intersections and fringes of the Venn diagram (Fig. 2). For depiction of global infection markers, the 40 most dominant genes in the intersection of all time points for both datasets each have been selected. For depiction of time point specific infection markers, the 10 most dominant genes of every gene set specific to each time point have been selected. The normalised read counts, averaged over replicate samples, were subjected to hierarchical clustering and are depicted per condition (GFP+, GFP- and Mock-infected) and along the time axis. Panels **A** and **B** show global infection markers of the early samples and the late samples, respectively. Genes in the intersection of global infection markers from both datasets are marked with an asterisk. Panels **C** and **D** show time point specific markers of the early samples and late samples, respectively. GFP-expression patterns were either intermediate between Mock-infected and GFP+ (**AC**) or more similar to Mock-infected (**BD**). Distinct time-courses are visible in **C** and **D**.

Although DEGA compared GFP⁺ and mock-infected samples, GFP⁻ cells included in clustering showed intermediate profiles during early infection (e.g., *gdcA*, DDB_G0284999, *cdk1* clusters), consistent with PCA results, but more closely resembled mock-infected controls at later stages. An exception was persistent intermediate regulation of *uduA2/A3*, possibly reflecting bystander responses to shed Mm envelope material, which is known to elicit transient G1/S delays (36) and may explain *cdk1* modulation in GFP⁻ cells.

These analyses define global and stage-specific transcriptional signatures of Mm infection, encompassing established defence genes (*zntB*) alongside novel candidates (*vacC*, *cdk1*) and poorly annotated genes, highlighting distinct transcriptional programs across infection stages.

### Autophagy and ESCRT pathways display pronounced temporal regulation during infection

Building on the global transcriptomic analysis, we next examined two pathways central to membrane homeostasis and pathogen control, ESCRT and autophagy (Fig. 4), using heatmaps of manually curated gene sets. In both datasets, GFP^+^ cells displayed distinct expression profiles from mock-infected and GFP⁻ populations, consistent with infection-induced remodelling of membrane repair and degradation pathways.

**Fig. 4.**
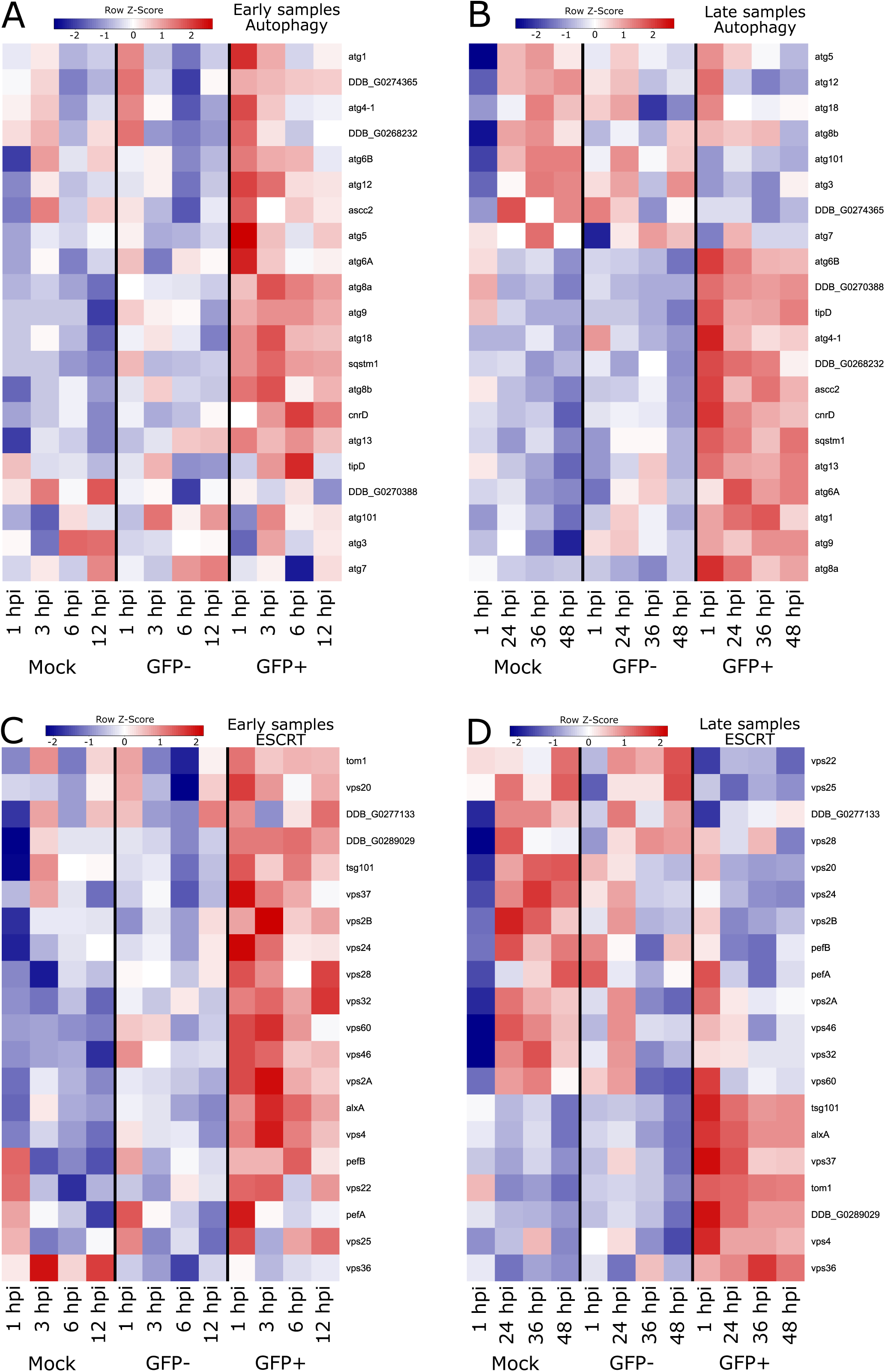
Autophagy and ESCRT related proteins are rewired in a time-dependent manner. Time-dependent regulation of selected genes related to autophagy (**A**, **B**) and ESCRT (**C**, **D**). The normalised read counts, averaged over replicate samples, were subjected to hierarchical clustering and are depicted per condition (GFP+, GFP- and Mock-infected) and along the time axis. Panels **A** and **C** show read counts for early samples, Panels **B** and **D** show read counts for late samples. Global upregulation of both machineries is visible, in particular a distinct succession of upregulated autophagy genes. Column legends at the bottom of panel **C** and **D** apply also for panels **A** and **B**.

The ESCRT pathway showed rapid and transient activation, with multiple components broadly induced in GFP⁺ cells at 1-3 hpi (Fig. 4A). These included *pefA/pefB*, which act upstream of ESCRT as sensors of membrane damage (37), followed by reduced expression at later time points relative to mock-infected and GFP^-^ cells. In contrast, core components (*tsg101, alxA, vps37, tom1, vps4,* and *vps36*) remained relatively stable throughout infection, consistent with sustained basal ESCRT activity.

Autophagy genes exhibited a more complex, multi-wave transcriptional response (Fig. 4B, C). An early wave at 1 hpi included *atg1, atg4, atg5, atg6A/B, atg12,* and *ascc2*, encoding components involved in autophagy initiation and phagophore biogenesis, including the Atg12-Atg5 conjugation system (38). A second wave at 3 hpi comprised *atg8a/b, atg9, atg18, atg101, atg3,* and *sqstm1/p62*, followed by a third cluster (*cnrD, atg13,* and *tipD*) peaking at 12 hpi. During late infection, several genes (*atg6A/B, atg4, atg13, atg1, atg9, ascc2, sqstm1/p62,* and *cnrD*) remained upregulated, with *atg9* showing the most sustained induction across the entire time course. In contrast, components of the Atg8-conjugation machinery (*atg7, atg3*) showed weaker and more variable regulation, while several autophagy genes (*atg5, atg12, atg18, atg101, atg3,* and *atg7*) became downregulated at later stages relative to mock-infected and GFP-cells, suggesting selective attenuation of the autophagy program during established infection.

Together, these data reveal distinct temporal regulation of ESCRT and autophagy during Mm infection. ESCRT components are rapidly induced during early infection, consistent with a role in sensing and repairing acute membrane damage, whereas autophagy displays a multi-phasic response with sustained regulation of selected components as infection progresses. The persistent induction of *atg9* is particularly notable and suggests a continued role in membrane trafficking and damage control beyond the early phase of infection. This sustained response, together with the broader temporal regulation of autophagy, motivated our subsequent functional analysis of Atg9.

### Pathway-level analysis reveals dynamic remodelling of membrane trafficking, proteostasis, and metabolism during Mm infection

To integrate the early and late datasets, we performed gene set enrichment analysis (GSEA) on KEGG pathways using LogFCs from each dataset separately (Fig. 5). At 1 hpi, many significant terms (p ≤ 0.05) were conserved between datasets, including endocytosis, autophagy, viral life cycle–HIV-1/ESCRT, and ribosome (Fig. 5A, B); notably, the HIV-1 term was driven largely by genes encoding ESCRT-associated proteins (*tsg101, vps32, alxA, vps4, vps20*). Proteasome-related terms were uniquely enriched at 1 hpi in the early dataset, while amino acid metabolism terms were enriched only in the late dataset (Fig. 5A, B).

**Fig. 5.**
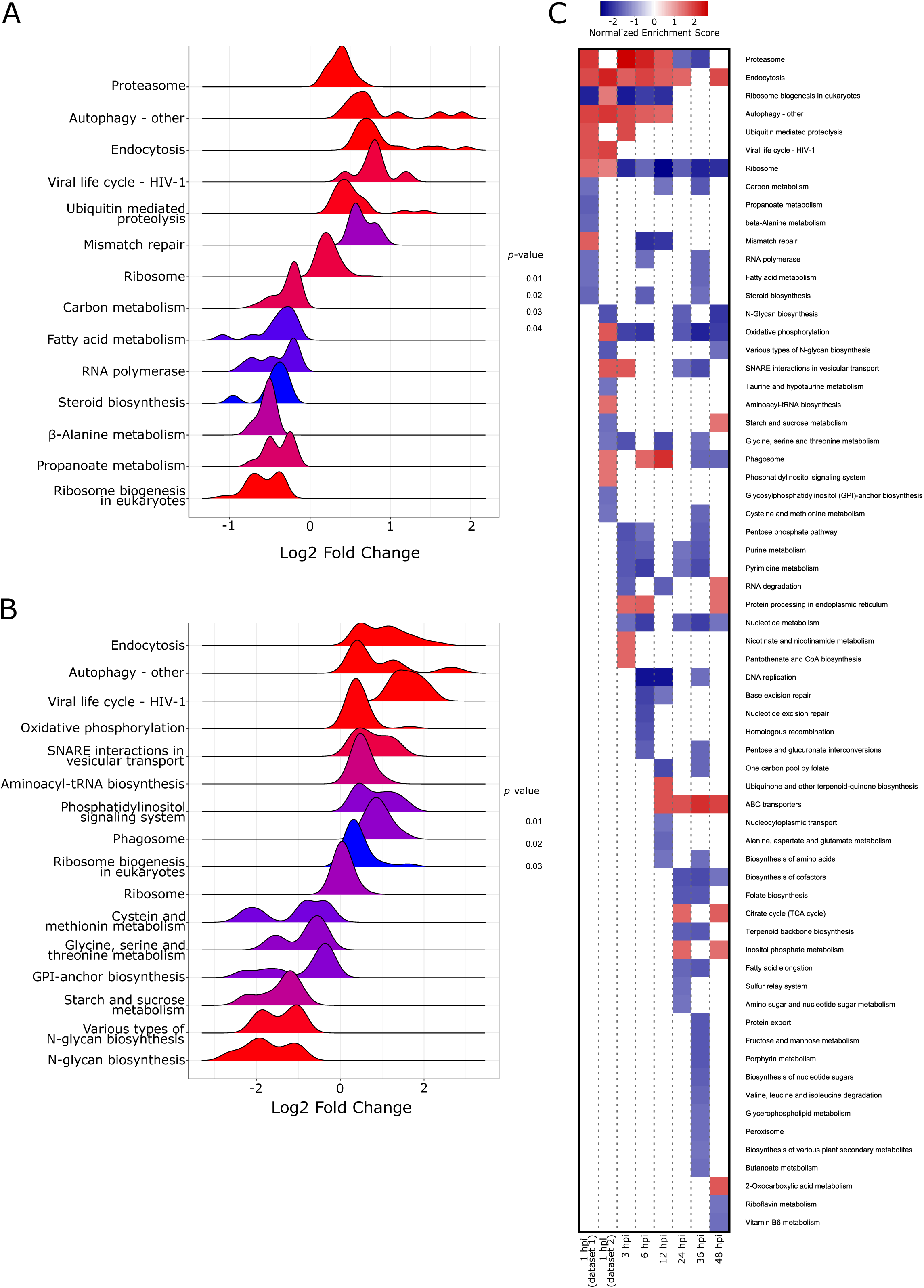
Merged pathway analysis. Comparison of datasets comprising early and late samples by pathway analysis. Gene set enrichment analysis (GSEA) was performed with the Kyoto Encyclopedia of Genes and Genomes (KEGG) annotation, significantly enriched terms (*p*-value ≤ 0.05) for the time point at 1 hpi from both datasets are shown in panels **A** and **B** as a ridgeplot. The graphs show the LogFC distributions within each term, the colour codes for *p*-value. In panel **B**, the term Glycosylphosphatidylinositol (GPI)-anchor biosynthesis has been abbreviated to GPI-anchor biosynthesis. The core enrichments of the term Viral life cycle - HIV-1 contain predominantly genes part of or associated with the ESCRT complex. Panel **C** shows the normalised enrichment score (NES) for every significantly enriched KEGG term at every time point in both datasets. A high NES, coded as red, indicates upregulation, a low NES, coded as blue, indicates downregulation of the genes within the respective term. In the column legend, dataset 1 and 2 refer to early and late samples, respectively.

Given this substantial overlap, we merged the analyses, plotting normalized enrichment scores for significantly enriched terms across all time points (Fig. 5C). Proteasome, endocytosis, and ribosome terms were enriched at nearly all time points: proteasome genes were upregulated early and downregulated from 24 hpi; endocytosis genes were upregulated throughout except at 36 hpi, partly overlapping with ESCRT genes but additionally including phagosome-maturation GTPases (*rab7A/B*, *rab8A/B*), previously linked to pathogen-containing vacuoles (4, 31), and ribosome/ribosome biogenesis genes were progressively downregulated or non-significant after the earliest time points, consistent with reduced biosynthetic activity and cell cycle progression. This potential signature of altered cell cycle progression is consistent with infection-dependent modulation of mTOR signalling (39) and cell cycle control. From 3 hpi, nucleotide metabolism, pentose phosphate pathway, and purine/pyrimidine metabolism genes were downregulated; from 6 hpi, DNA replication and base excision repair genes were downregulated or non-significant.

From 12 hpi onward, ABC transporter genes were consistently upregulated, including *abcG2* (linked to endosomal pH regulation (40)) alongside a*bcA3*, *abcG14*, *abcA2*, *abcB3*, *abcD2*, and *abcA6*. From 24 hpi, the TCA cycle was upregulated (non-significant at 36 hpi), while biosynthesis of cofactors was concurrently downregulated.

Overall, the data shows a dynamic reprogramming of host cellular functions throughout Mm infection, extending beyond classical antimicrobial pathways to encompass protein turnover, biosynthetic activity, metabolism, membrane trafficking, and cell cycle control. The progressive shift from early proteasomal and translational activity toward altered nucleotide metabolism, transporter expression, and central carbon metabolism reflects the changing cellular demands imposed by infection and the transition from vacuolar to cytosolic bacteria growth.

### The response of *D. discoideum* to *M. marinum* infection recapitulates signatures observed in human macrophages infected with Mtb

To assess conservation of these transcriptional responses in mammalian systems, we compared our datasets with five publicly available RNA-seq studies of Mtb-infected human macrophages (30, 41–44). Using human orthologs of Dd genes, we identified 1,351 DEGs shared across all datasets (Fig. 6A); the largest overlap (4,994 genes) occurred among studies using different Mtb/*M. bovis* strains, with 2,596 DEGs shared exclusively across Mtb-strain studies.

**Fig. 6.**
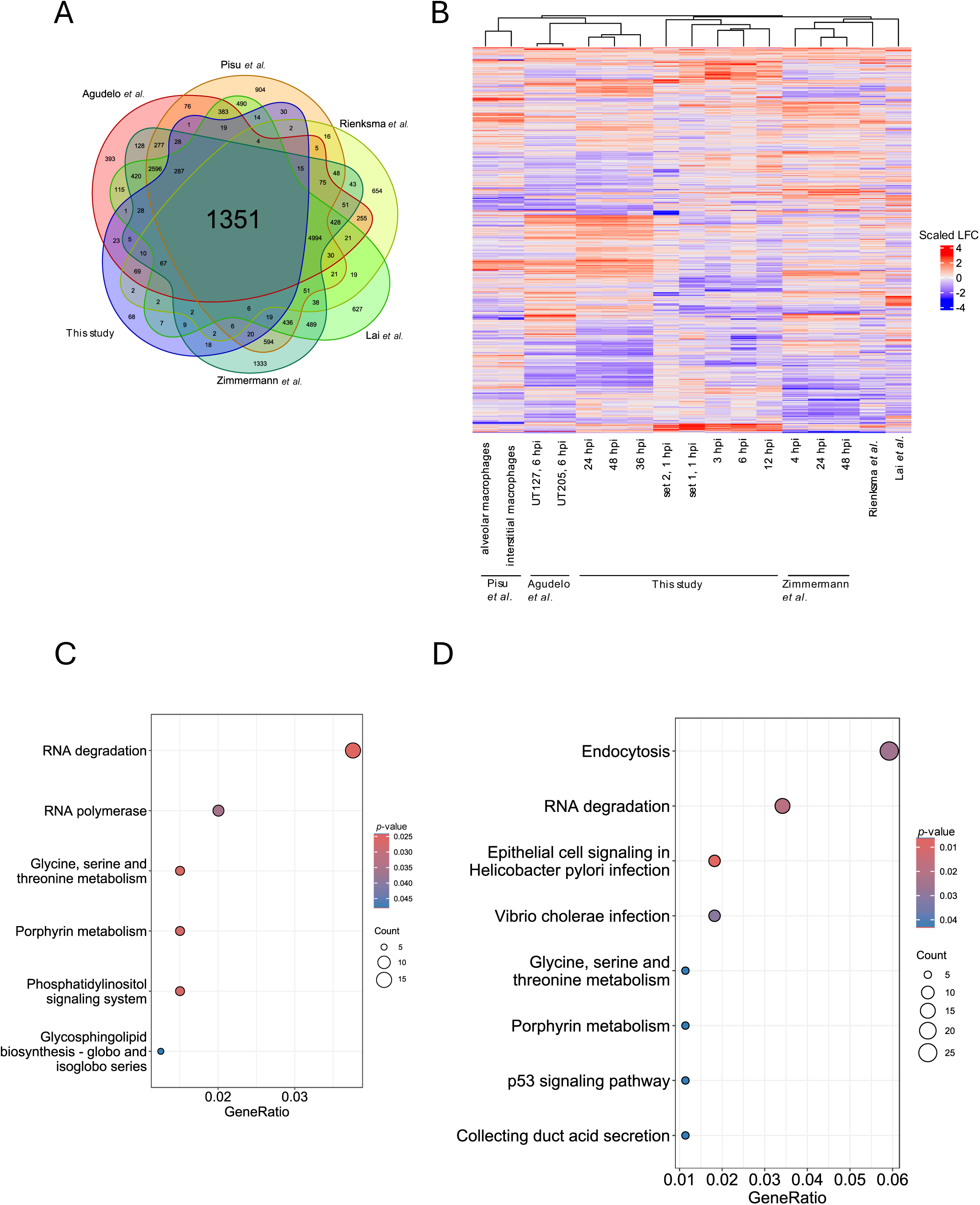
Important aspects of the host transcriptional signature are conserved in Mtb-infected human macrophages. Comparison of differentially expressed genes analysis (DEGA) of this study with five selected RNAseq studies on mycobacteria infection of human or mice macrophages. After mapping all gene identifiers to human orthologs, a set of 1351 genes was found to be shared with every of the five studies, in every condition (**A**). Hierarchical clustering of studies or, if present, different conditions within each study by scaled and centred log_2_ fold change (LogFC) is depicted as a clustered heatmap (**B**). High scaled LogFC is coded as red, a low scaled LogFC is coded as blue. High similarly between the time points at 24, 36 and 48 hpi from this study and the expression patterns observed by López-Agudelo et al. is visible. Genes with a consistent positive or negative LogFC in both sets, 24, 36 and 48 hpi from this study and human spleen macrophages infected with either the UT127 or UT205 Mtb strain were subjected to KEGG term enrichment of Dd gene identifiers (**C**) or human gene identifiers (**D**). The dotplots in **C** and **D** show significantly overrepresented KEGG terms (*p*-value ≤ 0.05), *p*-value is colour coded, the size of the genes enriched in each term is coded by dot size. The overrepresented terms are ordered by GeneRatio on the x-axis, which equals the number of queried genes over the number of genes associated with the respective KEGG term for the respective organism. Macrophages are abbreviated with Mps.

Hierarchical clustering of LogFCs for the 1,351 shared genes (Fig. 6B) separated our samples into early (1–12 hpi) and late (24–48 hpi) clusters. Notably, the late time points from our study clustered most closely with the dataset from López-Agudelo et al. (44), in which human spleen-derived macrophages were infected with two lineage 4 Mtb isolates (UT127 and UT205; MOI 10; 6 hpi), whereas the other macrophage datasets formed separate clusters. This similarity was further supported by moderate positive correlations between our late infection time points and both López-Agudelo conditions (Fig. S1). Among the 994 genes that could be compared across our late time points and both López-Agudelo conditions, 698 genes (70%) showed concordant LogFC direction, whereas 296 genes (30%) displayed discordant regulation. KEGG overrepresentation analysis of the 698 concordant genes (Fig. 6C, D) identified RNA degradation as conserved in both organisms, driven partly by Lsm proteins; in Dd, this included enrichment of RNA polymerase subunits (*rpa1, rpa2, rpc1, rpb8, rpb1, rpa5, rpb7, rpb3*). Glycine/serine/threonine metabolism was also conserved, corresponding to genes downregulated from 1 hpi onward (Fig. 5C, Table S3). In human macrophages, conserved enrichment included endocytosis, with *rab5c* and several ESCRT components (CHMP1A, CHMP1B, VPS25, VPS37A, and SNX1); the Dd SNX1 homologue, SnxA, has been implicated in phosphoinositide-dependent endosomal trafficking (45). Pathways related to acid secretion and *Vibrio cholerae* infection were also enriched, driven in part by V-ATPase subunits involved in vesicular and phagosomal acidification. Notably, the KEGG autophagy term revealed that *atg9* was upregulated in both Dd, and human macrophages infected with Mtb, with this response extending to later stages of infection. Among the autophagy-associated genes identified in this analysis, *atg9* was the only one showing a concordant transcriptional pattern across both systems together with sustained upregulation at later time points (Table S3). This conserved and temporally sustained regulation of *atg9* highlighted Atg9 as a candidate host factor for functional investigation.

Together, these analyses identify conserved transcriptional responses to mycobacterial infection involving membrane trafficking, ESCRT function, RNA metabolism, and phagosomal acidification, further supporting the relevance of the Dd-Mm model for investigating evolutionarily conserved host responses and providing a rationale for characterizing the role of Atg9 in mycobacterial infection.

### Atg9 is induced and recruited to the MCV to restrict intracellular growth

To functionally characterize the role of Atg9 during Mm infection, we first examined the transcriptional dynamics of *atg9*. Consistent with a role for Atg9 during infection, *atg9* expression was dynamically regulated throughout early and late infection stages (Fig. 7A and B). Infected GFP⁺ cells displayed a distinct temporal transcriptional profile compared to both GFP- and mock-infected conditions, indicating that *atg9* induction is specifically associated with infection. At the protein level, immunoblot analysis confirmed an increase in Atg9 abundance across time points, with a significant upregulation observed at 6 hpi (Fig. 7C and D), supporting our findings from transcriptional analysis. To assess the spatial dynamics of Atg9 during infection, live-cell imaging of Dd expressing Atg9-mCherry infected with GFP-expressing Mm was performed (Fig. 7E). Atg9-positive structures dynamically associated with and trafficked in close proximity to the MCV throughout infection. This was further examined at higher throughput by high-content imaging (Fig. 7F), which revealed markedly increased recruitment and accumulation of Atg9-positive structures around MCVs containing virulent Mm compared to the ΔRD1 mutant. Quantification confirmed a significantly higher proportion of Atg9-positive compartments during infection with the wt strain compared to ΔRD1 (Fig. 7G), indicating that Atg9 recruitment scales with bacterial virulence and the extent of MCV membrane damage.

**Fig. 7.**
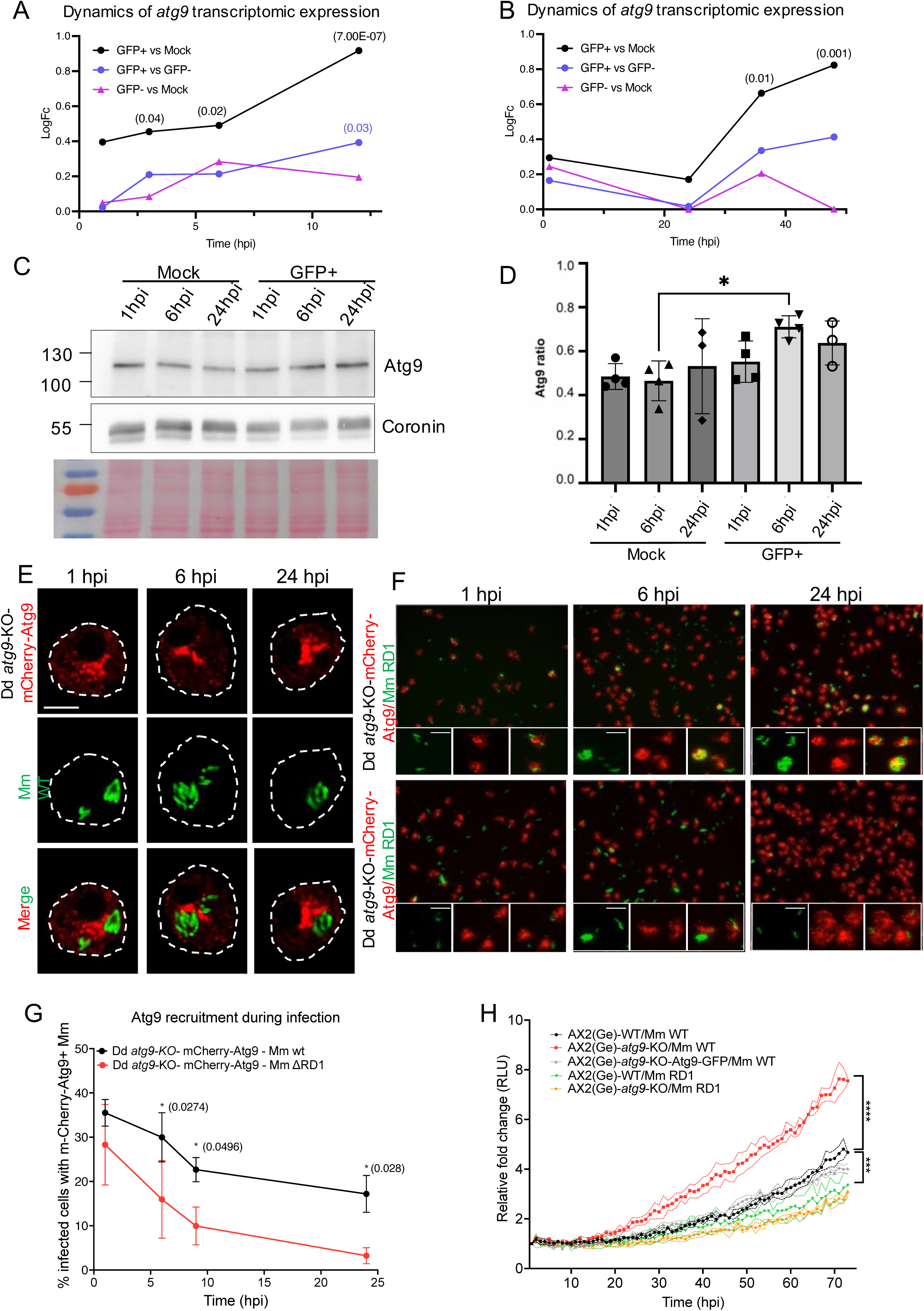
Temporal regulation, recruitment, and functional role of Atg9 during Mm infection in Dd. **(A, B)** Time-resolved transcriptomic profiling of *atg9* expression during early (A) and late (B) stages of infection. LogFC of *atg9* transcript levels was measured over the course of infection in infected (GFP⁺), bystander (GFP⁻), and mock-infected cells, showing dynamic infection-associated transcriptional regulation. Comparisons are shown between GFP⁺ versus mock-infected, GFP⁺ versus GFP⁻, and GFP⁻ versus mock-infected conditions. **(C)** Representative immunoblot analysis of Atg9 protein levels in mock-infected and Mm-infected cells collected at 1, 6, and 24 hours post-infection (hpi). Coronin was used as loading control. **(D)** Quantification of Atg9 protein abundance normalized to Coronin. Atg9 levels increased during infection, with a significant accumulation observed at 6 hpi compared to mock-infected controls. Data are presented as normalized Atg9/Coronin ratios. **(E)** Representative live images of *atg9*-KO expressing Atg9-mCherry cell infected with Mm wt expressing GFP; scale bars, 5 μm. Images were collected and processed using Leica Stellaris microscope with lightning default processing. **(F)** Representative fluorescence microscopy images of Dd *atg9*-KO expressing Atg9-mCherry infected with GFP-expressing Mm wt or ΔRD1 at indicated time points. Atg9 recruitment to the MCV is visualized by mCherry signal accumulation surrounding intracellular bacteria; scale bars, 10 μm. **(G)** Quantification of Atg9-positive MCVs during infection with Mm wt or ΔRD1. The percentage of infected cells displaying Atg9 recruitment was determined at indicated time points. **(H)** Intracellular growth of Mm wt and ΔRD1 in Dd wt, *atg9*-KO, and complemented strains measured by relative luminescence units (RLU) over time. wt Mm displayed enhanced intracellular replication in *atg9* knockout cells compared to wt host cells, whereas complementation restored bacteria growth control to wt levels. No significant differences were observed during infection with the ΔRD1 mutant. Data represent mean ± SEM from at least three independent experiments. Statistical significance was determined using 2way ANOVA; *P* < 0.05, P < 0.01, *P* < 0.001.

Finally, functional analysis demonstrated that loss of Atg9 enhances intracellular bacteria growth. Lux-expressing Mm wt exhibited significantly increased growth in *atg9*-KO cells compared to wt Dd, whereas complementation of the knockout restored bacteria growth to wt levels (Fig. 7H). In contrast, infection with the ΔRD1 mutant resulted in comparable intracellular bacteria growth in wt and *atg9*-KO cells, indicating that the Atg9-dependent restriction mechanism specifically impacts Mm strains capable of accessing the cytosolic compartment.

### Atg9 preserves MCV integrity and restricts early Mm escape to the cytosol

To determine whether Atg9 contributes to membrane damage responses and maintenance of MCV integrity, we first assessed ESCRT-mediated membrane repair dynamics following chemically induced lysosomal damage using LLOMe treatment. Live-cell imaging of GFP-Vps32-expressing Dd cells revealed rapid formation of GFP-Vps32-positive puncta following membrane damage in both wt and *atg9*-KO cells (Fig. 8A). Quantification of responding cells showed similar kinetics of GFP-Vps32 recruitment in the absence of Atg9 (Fig. 8B), while analysis of total GFP-Vps32-positive area demonstrated reduced accumulation of GFP-Vps32 structures in *atg9*-KO cells compared to wt controls (Fig. 8C). Consistently, area under the curve analysis confirmed a significantly reduced cumulative GFP-Vps32 response in *atg9*-KO (Fig. 8D), indicating that while initial ESCRT recruitment is intact, the magnitude and efficiency of the subsequent repair response are reduced in the absence of Atg9.

**Fig. 8.**
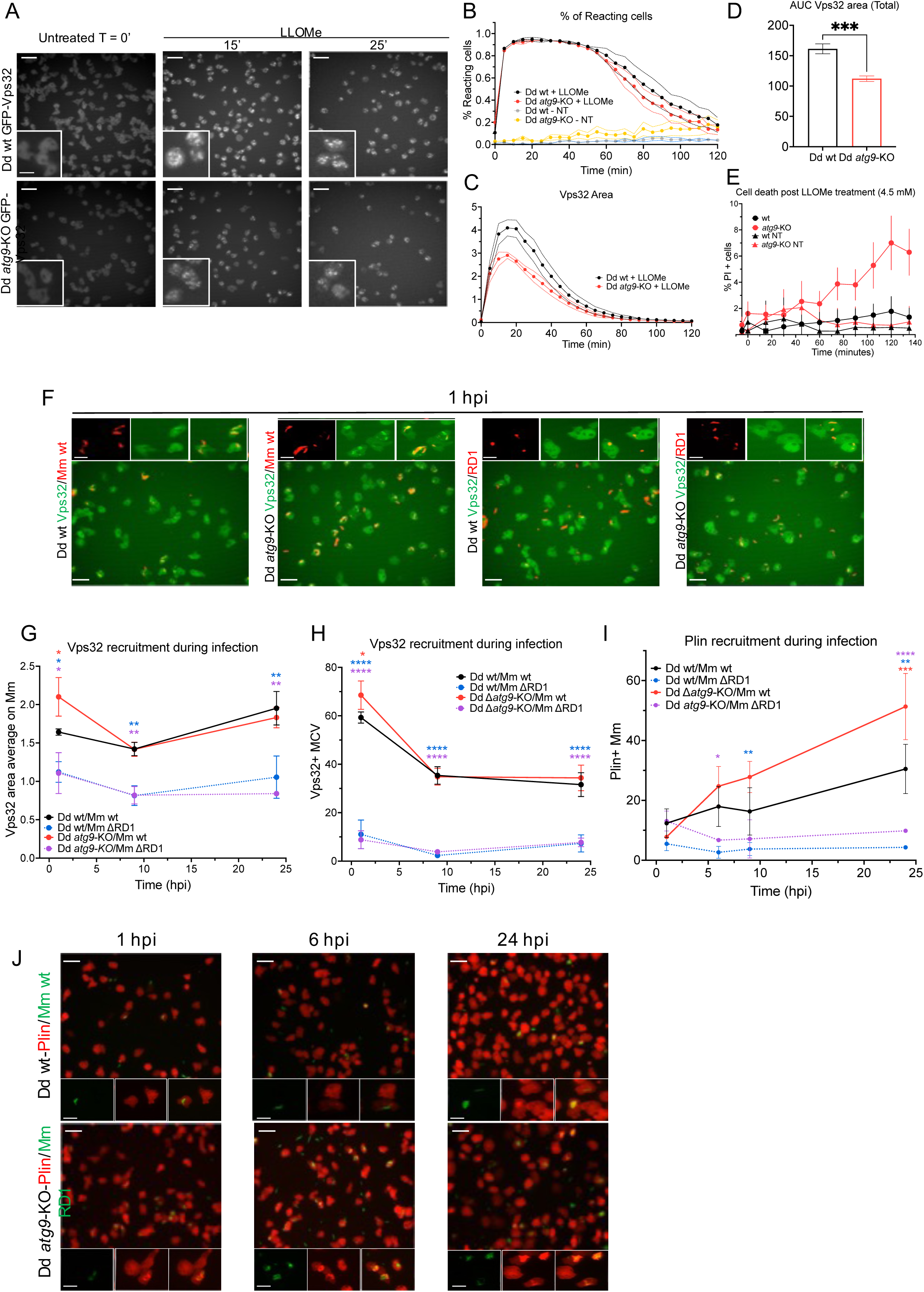
Atg9 promotes membrane damage responses and restricts early Mm escape to the cytosol. **(A)** Representative live-cell fluorescence microscopy images of Dd wt and *atg9*-KO cells expressing GFP-Vps32 following LLOMe treatment at the indicated time points. Insets highlight GFP-Vps32-positive puncta formed in response to chemically induced endolysosomal membrane damage; scale bars, 10 μm. **(B)** Quantification of the percentage of responding cells, defined by the presence of detectable GFP-Vps32-positive puncta, following LLOMe treatment. *atg9*-deficient cells exhibited altered kinetics of GFP-Vps32 recruitment compared to wild-type cells. **(C)** Quantification of total GFP-Vps32-positive area per cell over time following LLOMe treatment. The temporal dynamics of GFP-Vps32 accumulation reveal differences in membrane damage response between wt and *atg9*-KO cells. **(D)** AUC analysis of GFP-Vps32-positive area shown in panel C, providing an integrated measure of the cumulative membrane repair response. Significant differences indicate impaired regulation of membrane damage sensing and/or repair in the absence of Atg9. **(E)** Quantification of PI-positive cells over time following treatment with 4.5 mM LLOMe in wt and *atg9*-KO cells, alongside untreated (NT) controls for each genotype. Data are presented as mean ± SEM. **(F)** Representative fluorescence microscopy images of Dd wt and *atg9*-KO cells expressing GFP-Vps32 during infection with Mm wt or ΔRD1. Insets highlight GFP-Vps32 recruitment to MCVs; scale bars, 10 μm. (**G)** Quantification of GFP-Vps32 area average on Mm during infection. Infection with Mm wt induced significantly increased GFP-Vps32 recruitment in *atg9*-KO cells compared to wt controls, whereas ΔRD1 infection elicited minimal recruitment in both host backgrounds. **(H)** Quantification of GFP-Vps32-positive MCVs during infection. Infection with Mm wt induced significantly increased GFP-Vps32 recruitment in *atg9*-KO cells compared to wt controls, whereas ΔRD1 infection elicited minimal recruitment in both host backgrounds. **(I)** Quantification of cytosolic accessibility over time. Mm wt gained access to the host cytosol significantly earlier and more frequently in *atg9*-KO cells, whereas ΔRD1 bacteria remained largely vacuole-confined regardless of host genotype. **(J)** Representative fluorescence microscopy images of wt and *atg9*-KO cells expressing the cytosolic accessibility reporter mCherry-Plin during infection with Mm WT or ΔRD1. Insets show magnified views of representative intracellular bacteria; scale bars, 10 μm.

**Fig. 9.**
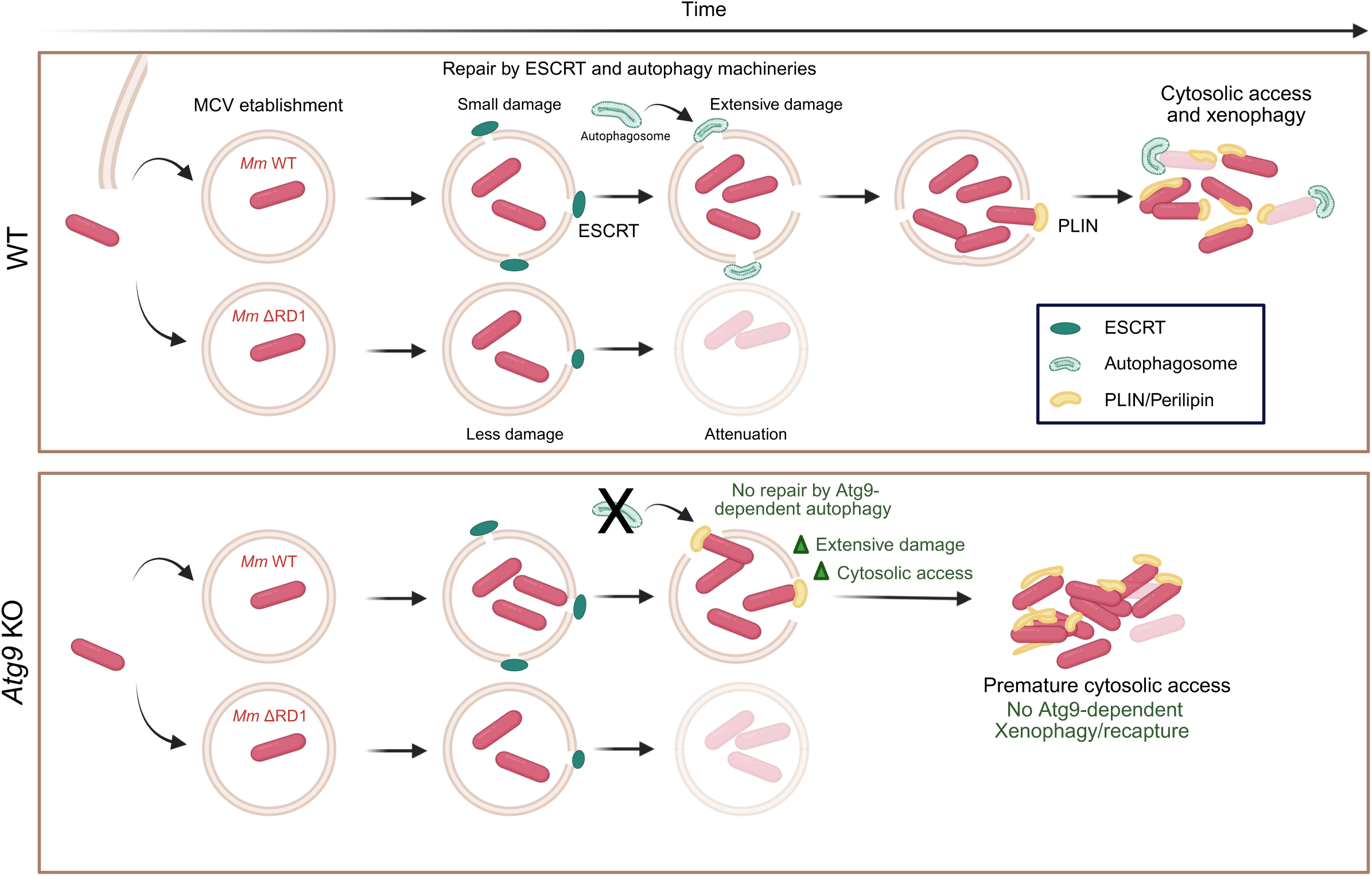
Disruption of Atg9-dependent autophagy accelerates Mm escape to the cytosol and enhances intracellular proliferation. Atg9 sustains ESCRT-mediated membrane repair to preserve Mm vacuolar confinement. In wt cells, ESX-1-dependent damage to the MCV triggers sequential recruitment of ESCRT machinery and Atg9-dependent autophagy to repair membrane lesions and delay bacteria escape. Once repair capacity is exceeded, cytosolic bacteria are targeted by xenophagy to limit intracellular proliferation. The ESX-1-deficient ΔRD1 mutant induces limited membrane damage that is efficiently resolved by ESCRT-mediated repair, maintaining stable vacuolar confinement. In *atg9*-deficient cells, ESCRT recruitment remains intact, but defective autophagy-dependent reinforcement of membrane repair results in unresolved damage, premature escape of Mm wt to the cytosol, and increased intracellular replication. Thus, Atg9 couples membrane repair with xenophagy to restrict pathogen access to the cytosolic replicative niche.

To assess whether this impaired repair response translates into a functional consequence for membrane integrity, we monitored cell death following LLOMe-induced damage using propidium iodide (PI) uptake as a readout, given that PI is excluded by intact plasma membranes and only labels cells that have lost integrity. Quantification over time revealed a progressive increase in the proportion of PI-positive *atg9*-KO cells following LLOMe treatment at 4.5 mM, exceeding the levels observed in wt cells, whereas untreated cells of both cell lines displayed minimal cell death throughout the imaging period (Fig. 8E; Fig. S2A). To determine whether this phenotype scaled with the severity of membrane damage, we repeated the assay using a higher LLOMe concentration (9 mM) (Fig. S2B). Quantification confirmed that cell death was overall increased at the higher dose in both cell lines, and the difference between wt and *atg9*-KO cells was further exacerbated, with *atg9*-KO cells reaching markedly higher levels of PI positivity than wt cells compared to the 4.5 mM condition (Fig. S2C). Together, these results indicate that loss of Atg9 compromises the efficiency of membrane damage repair, resulting in increased susceptibility to membrane-damage-induced cell death, an effect that becomes more pronounced with increasing severity of membrane damage.

We next investigated whether this altered membrane repair capacity impacts MCV integrity during Mm infection. Representative live-cell imaging revealed recruitment of GFP-Vps32 to MCVs in cells infected with virulent Mm wt, whereas recruitment remained minimal during infection with the ΔRD1 mutant (Fig. 8F, Fig. S2D), confirming that ESCRT activation depends on ESX-1-mediated membrane disruption. Consistent with the intact initial damage sensing observed following LLOMe treatment, the proportion of GFP-Vps32-positive MCVs was comparable between wt and *atg9*-KO cells, confirming that loss of Atg9 does not impair initial recruitment of ESCRT machinery to damaged MCV (Fig. 8F). However, *atg9*-KO cells displayed a significantly increased GFP-Vps32-positive area associated with MCVs (Fig. 8G). Rather than reflecting a stronger repair response, this enlarged GFP-Vps32 signal is consistent with a reduced overall repair efficiency, in which cells try to compensate defects in Atg8ylation-dependent membrane repair by persistent or repeated ESCRT engagement, which fails to fully resolve membrane damage, allowing it to accumulate over the course of infection.

To directly assess bacteria access to the host cytosol, intracellular bacteria were visualized using the cytosolic accessibility reporter mCherry-Plin (Fig. 8I,J; Fig. S2E). Upon cytosolic exposure, the perilipin ortholog partitions to the hydrophobic surface of the Mm envelope, providing a readout of bacteria cytosolic accessibility. Mm wt progressively acquired cytosolic accessibility in wt cells, whereas this process was significantly accelerated in *atg9*-KO cells, with a higher proportion of mCherry-Plin-positive bacteria at all time points, consistent with earlier escape from the MCV (Fig. 8I). This phenotype is consistent with impaired membrane repair, whereby sustained damage caused by growing Mm may exceed the repair capacity of Atg9-deficient cells, promoting loss of vacuolar confinement. In contrast, the ΔRD1 mutant remained largely inaccessible to the cytosolic reporter in both host backgrounds, consistent with persistent vacuolar confinement and the established ESX-1 dependence of Mm escape to the cytosol through ESX-1-mediated membrane damage (14, 15, 46).

Together, these findings demonstrate that Atg9 is dispensable for initial sensing and recruitment of the ESCRT machinery to damaged membranes, but is required for efficient resolution of that damage and preservation of MCV integrity during the earliest stages of infection. Loss of Atg9 leads to reduced membrane repair efficiency, enhanced ESX-1-dependent vacuolar damage, and accelerated Mm escape to the cytosol, thereby promoting the transition from a vacuolar to cytosolic lifestyle that supports intracellular bacteria proliferation. Mm then grows unrestricted in the cytosol in absence of Atg9 because of defective xenophagy, underscoring that Atg9’s role extends beyond membrane repair to the subsequent xenophagic capture of bacteria that do escape to the cytosol.

## DISCUSSION

Using an unprecedented temporal resolution to interrogate this model system, we defined global and time-resolved transcriptional signatures associated with Mm infection in Dd. Comparison with transcriptomic data from human spleen-derived macrophages infected with South American Mtb isolates revealed substantial concordance in transcriptional responses, including several conserved genes and biological pathways.

Among the most strongly regulated genes, *IliL* and *IliA*, encoding putative glycoproteins previously reported to be induced following both Lp and Mm infection of Dd, and *rigA*, a component of the *rasD* signalling network involved in cell-fate regulation during slug-stage differentiation, showed particularly pronounced changes. This is consistent with previous evidence that *rasD* is upregulated during Mm infection (31). Conversely, members of the *uduA1/A2/A3*, *uduB*, and *uduG* families, previously associated with the MAPK regulator DupA during Lp infection, exhibited strong negative LogFCs, suggesting differential regulation of these pathways during mycobacteria infection.

Small heat-shock proteins of the HspG family were among the most consistently upregulated genes, despite cells being maintained at a temperature close to the upper range compatible with Mm growth. This contrasts with the HspG downregulation previously reported during Mm infection of Dd (33, 34), highlighting potential differences in the regulation of the heat-shock response depending on experimental conditions.

ESCRT and autophagy-related genes displayed distinct temporal expression patterns, particularly within the autophagy machinery components. Subunits of the Atg1 complex and Atg12 conjugation system were induced early, whereas Atg7 and Atg3, which participate in Atg8 lipidation, showed comparatively limited early induction. Notably, *atg8* itself was upregulated, potentially reflecting functions beyond canonical autophagy, including its reported involvement in ESCRT-dependent membrane repair (21). These distinct transcriptional dynamics suggest that individual autophagy components may be differentially regulated and recruited to fulfil specialized functions during Mm infection. The ESCRT machinery is well established as being involved in repair of small membrane damages and was observed to be upregulated from early time points on with sustained upregulation also at late infection time points. Recently, ESCRT recruitment was found to be regulated by TrafE, which additionally acts at the intersection of ESCRT and autophagy recruitment (16). Together with the findings in this study, this strongly supports further research into dissection of time-dependent expression and recruitment of TrafE-dependent components of the ESCRT and autophagy machinery.

Kjellin et al. profiled Dd infected with Mm at a single early time point (2.5 hpi). Our time-resolved analysis recapitulated several of their major transcriptional signatures, including pathways associated with phagosome maturation and acidification, as well as induction of ESCRT- and autophagy-related machineries. In contrast, ABC transporter genes, which were reported to be downregulated at 2.5 hpi, were predominantly upregulated at later stages in our dataset, highlighting the temporal dependence of the host transcriptional response. We also identified signatures consistent with altered cell-cycle regulation (36), coinciding with sustained downregulation of nucleotide metabolism from 3 hpi onward (Fig. 5C).

Comparison of LogFC profiles with published human and mouse macrophage–*Mtb* infection datasets revealed the strongest cross-species concordance between our late infection time points (24–48 hpi) and human spleen-derived macrophages from Agudelo *et al.* Notably, the correlation between the Dd-Mm and Agudelo *et al.* datasets was higher than that observed between any pair of the published Mtb-infected macrophage datasets, indicating a particularly strong transcriptional similarity between the Dd-Mm and human macrophage infection systems despite their distinct host and pathogen species. To our knowledge, this represents one of the most extensive quantitative comparisons of transcriptional responses between the Dd-Mm and mammalian macrophage-Mtb infection models.

Overall, our findings strongly support the study of mycobacteria infection of Dd as a model for the human innate immune system but in absence of intercellular cytokine signalling (34). These findings extend previous Dd transcriptomic studies by resolving the host response across infection stages and identifying both conserved and temporally restricted signatures. The pronounced regulation of ESCRT and autophagy pathways further supports membrane damage, repair, and bacteria killing and degradation as key components of the host response, while the temporal regulation of cell-cycle and heat-shock pathways highlights additional processes that warrant further investigation. Future dual RNA-seq analyses incorporating the bacterial transcriptome could provide a more integrated view of host–pathogen dynamics and clarify how temporal changes in host pathways relate to bacterial adaptation and intracellular proliferation, with potential implications for understanding mycobacterial pathogenesis and identifying host-directed therapeutic strategies.

Following up on the transcriptomic results, we identify Atg9 as a critical host restriction factor that limits Mm intracellular proliferation by preserving MCV integrity and preventing bacteria access to the cytosol. Temporal transcriptomic and protein analyses revealed dynamic induction of Atg9 during infection, with expression specifically enriched in infected cells. This induction was accompanied by robust recruitment of Atg9 to the MCV, particularly during infection with virulent Mm wt, linking its localization to ESX-1-dependent membrane damage. Consistent with this, loss of Atg9 selectively enhanced intracellular replication of Mm wt, with no effect on the attenuated ΔRD1 mutant, indicating that Atg9-mediated restriction specifically counteracts virulence-associated membrane damage and phenocopies the Mm growth phenotype in *atg1*-KO cells for both wt and ΔRD1 infections (39).

Loss of Atg9 did not impair initial ESCRT recruitment but compromised subsequent repair efficiency. Following LLOMe treatment, *atg9*-KO cells recruited GFP-Vps32 with kinetics comparable to wt cells but accumulated significantly less GFP-Vps32 signal over time, indicating that Atg9 acts downstream of damage sensing to support productive ESCRT-mediated repair. During Mm infection, by contrast, *atg9*-KO cells displayed enlarged, unresolved GFP-Vps32 assemblies and accelerated escape to the cytosol. This difference likely reflects the distinct dynamics of membrane damage: LLOMe induces acute and largely self-limited lysosomal permeabilization, whereas Mm causes localized, sustained, and progressive ESX-1-and PDIM-dependent damage to the MCV (46). Thus, impaired repair in *atg9*-KO cells may limit the overall accumulation of Vps32 following transient LLOMe-induced damage, whereas persistent Mm-induced damage may drive prolonged ESCRT-III recruitment, resulting in enlarged and unresolved assemblies. This phenotype resembles that observed in E3 ubiquitin ligase *trafE*-KO cells, where ESCRT-III accumulates without productive Vps4-dependent disassembly, providing a precedent for persistent ESCRT assemblies as a signature of defective membrane repair (16).

Together, these findings indicate that Atg9 is dispensable for initial damage sensing and ESCRT recruitment but is required for efficient completion of membrane repair, particularly under sustained or recurrent damage such as that generated during active Mm infection. This places Atg9 at the interface between autophagy and membrane repair, extending its role beyond canonical degradative autophagy and supporting emerging models of Atg9 in lysosomal repair and non-canonical membrane remodelling (29, 47). It is also consistent with the membrane ATG8ylation framework, in which autophagy-related proteins participate in broader responses to pathogen-induced membrane stress beyond degradative autophagy (20, 48). Together with the ESX-1-dependent recruitment of Atg9 to damaged MCVs, the increased membrane damage and accelerated bacteria escape observed in its absence support a model in which Atg9-dependent autophagic machinery functionally cooperates with ESCRT-mediated repair to maintain MCV integrity. This coordinated response ultimately limits Mm access to the cytosolic replicative niche and contributes to host defence against ESX-1-mediated pathogenesis.

## MATERIAL AND METHODS

### *D. discoideum* and mycobacteria strains, culture, and plasmids

Dd and Mm strains and plasmids are listed in Table 1. Dd Ax2(Ka) was cultured axenically at 22°C in HL5c medium (Formedium) supplemented with penicillin (100 U/mL) and streptomycin (100 μg/mL). Plasmids were introduced by electroporation and selected with the appropriate antibiotic; hygromycin was used at 50 μg/mL for reporters integrated at the safe-haven *act5* locus. Mm (M strain) wt was cultured in 7H9 medium supplemented with 0.2% glycerol, 0.05% Tween-20, and 10% OADC. GFP-expressing bacteria were generated by transformation with the *msp12::GFP* vector and maintained with 50 μg/mL kanamycin.

**Table 1.**
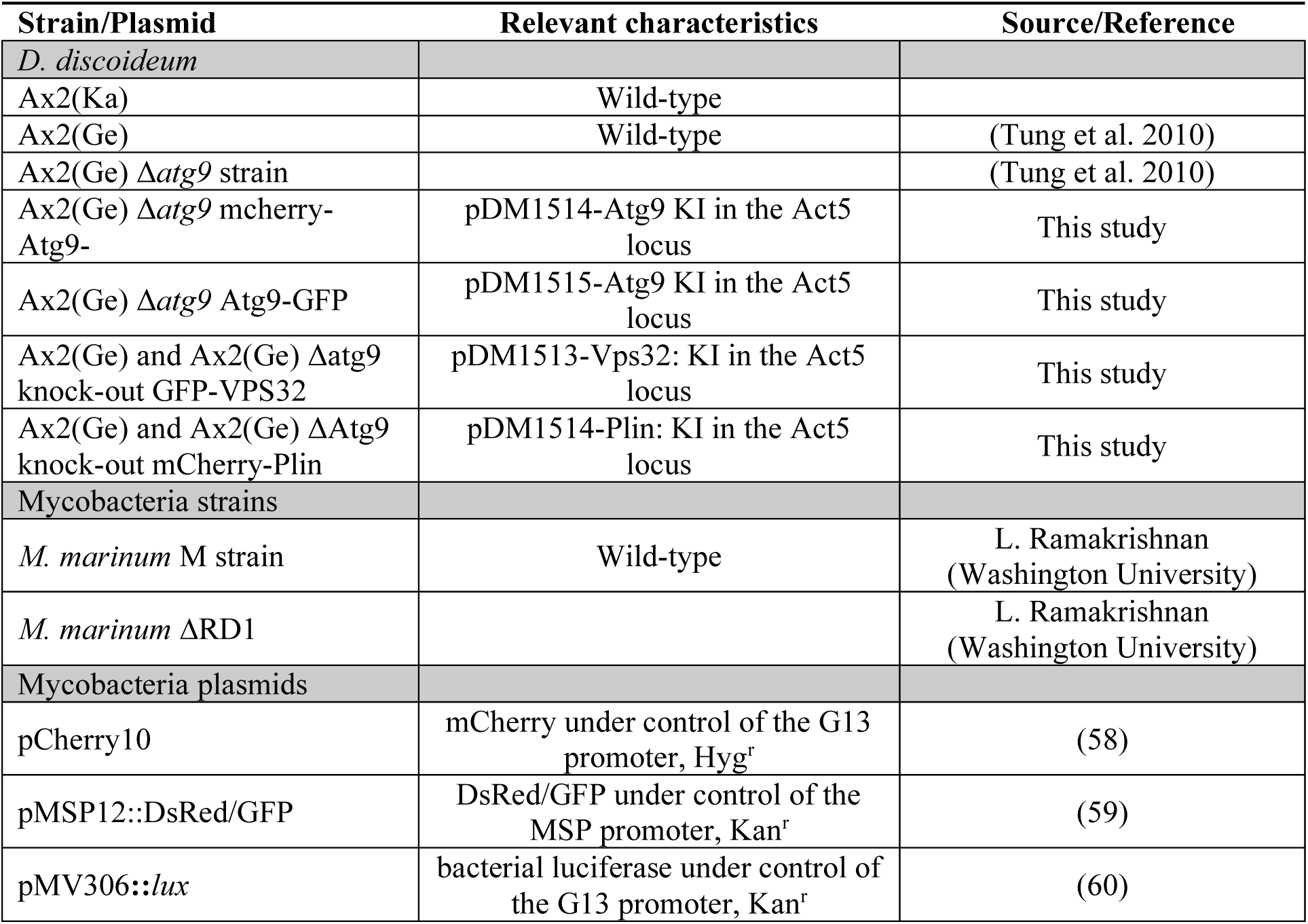
List of *D. discoideum* and *M. marinum* strains used in this study.

| Strain/Plasmid | Relevant characteristics | Source/Reference |
| --- | --- | --- |
| *D. discoideum* |  |  |
| Ax2(Ka) | Wild-type |  |
| Ax2(Ge) | Wild-type | (Tung et al. 2010) |
| Ax2(Ge) $\Delta atg9$ strain | | (Tung et al. 2010) |
| Ax2(Ge) $\Delta atg9$ mcherry-Atg9- | pDM1514-Atg9 KI in the Act5 locus | This study |
| Ax2(Ge) $\Delta atg9$ Atg9-GFP | pDM1515-Atg9 KI in the Act5 locus | This study |
| Ax2(Ge) and Ax2(Ge) $\Delta atg9$ knock-out GFP-VPS32 | pDM1513-Vps32: KI in the Act5 locus | This study |
| Ax2(Ge) and Ax2(Ge) $\Delta Atg9$ knock-out mCherry-Plin | pDM1514-Plin: KI in the Act5 locus | This study |
| *Mycobacteria* strains |  |  |
| *M. marinum* M strain | Wild-type | L. Ramakrishnan (Washington University) |
| *M. marinum* $\Delta RD1$ | | L. Ramakrishnan (Washington University) |
| *Mycobacteria* plasmids |  |  |
| pCherry10 | mCherry under control of the G13 promoter, Hyg<sup>r</sup> | (58) |
| pMSP12::DsRed/GFP | DsRed/GFP under control of the MSP promoter, Kan<sup>r</sup> | (59) |
| pMV306::lux | bacterial luciferase under control of the G13 promoter, Kan<sup>r</sup> | (60) |

### Infection assay

Infections were performed as previously described (49). Briefly, overnight-grown GFP-expressing *M. marinum* were centrifuged, resuspended in HL5c, and spinoculated onto adherent *D. discoideum* cells at an MOI of 25. Following removal of uningested bacteria, infected cells were resuspended in filtered HL5c at 1 × 10⁶ cells/mL and incubated at 25°C with shaking at 130 rpm. Penicillin (5 U/mL) and streptomycin (5 μg/mL) were added to suppress extracellular bacteria growth. Samples were collected in two independent time-course series covering early (1, 3, 6, and 12 hpi) and late (1, 24, 36, and 48 hpi) infection stages. A non-infected, mock-infected control was included at each time point to account for transcriptional changes associated with culture conditions and treatment.

### Flow cytometry and fluorescence-activated cell sorting (FACS)

To enrich for host and bacterial RNA and obtain defined infection-state populations, infected (GFP⁺) and uninfected bystander (GFP⁻) *D. discoideum* cells were separated by FACS. Mm-GFP-infected and mock-infected cells were collected, centrifuged (5 min, 1500 rpm), and resuspended in HL5c. Cells were sorted on an Astrios cell sorter (Beckman) at 4°C, with both the input holder and collection rack cooled. Cells were first gated by size (forward scatter) and granularity (side scatter), followed by separation of GFP⁺ and GFP⁻ populations based on GFP fluorescence (FITC) relative to autofluorescence (PE). Conservative gates were used to minimize cross-contamination, with identical settings applied to mock-infected controls. Approximately 5 × 10⁵ cells per fraction were collected, centrifuged, and resuspended in TRI Reagent (T9424, Sigma-Aldrich) for RNA extraction (50).

### RNA isolation, library preparation, and sequencing

Total RNA was extracted using the Direct-zol RNA Extraction Kit (Zymo Research) and treated with DNase I (0.25 U/μg RNA, 15 min, 25°C). RNA concentration and integrity were assessed using Qubit 4.0 (Invitrogen) and an Agilent 2100 Bioanalyzer. Libraries were prepared from 100 ng RNA using the Ovation Universal System (NuGEN), with double-sided bead size selection, end repair, barcoded adapter ligation, strand selection, and custom oligonucleotide-mediated depletion of Dd and Mm rRNAs. Libraries were amplified by 18 PCR cycles, quality-controlled using an Agilent TapeStation, pooled in six-plex at approximately equimolar concentrations, and sequenced as 50-bp single-end reads on an Illumina HiSeq 4000.

### RNA-seq mapping and differential expression analysis

50-bp single-end reads were aligned to the Dd genome (51, 52) using TopHat v2.0.13 with Bowtie2 v2.2.4, allowing only uniquely mapped reads (--max-multi-hits 1). Gene-level counts were generated with HTSeq v0.6.1 using the DictyBase GFF annotation (February 2019; -t exon --stranded=yes -m union). Counts were analysed in R v4.0.3 using DESeq2 v1.30.1 (Love et al., 2014) with HTS filtering, after removal of residual rRNA-derived counts. GFP⁺, GFP⁻, and mock-infected samples were compared pairwise at each time point. Fold changes were estimated using the *ashr* method (53), and *p*-values were adjusted using the Benjamini– Hochberg procedure, with FDR ≤ 0.05 and |log_2_ fold change| ≥ 0.585 (1.5-fold) used to define differentially expressed genes. For heatmap visualization, genes were ranked by a dominance score calculated as absolute (LogFC) × −log10(FDR), with the maximum score across time points used for gene selection.

### Heatmaps of normalized counts

Gene counts were filtered for residual ribosomal genes and low-expression genes using filterByExpr (edgeR) (54), normalized using variance-stabilizing transformation, and batch-corrected for experimental date using limma. Heatmaps were generated with heatmap.2 (gplots v3.1.1), with data scaled and centered by gene.

### Pathway analysis

Pathway enrichment was performed using clusterProfiler and KEGG annotations (accessed 27 January 2023)(55). For GSEA, genes were ranked by LogFC and analysed using 10,000 permutations, with significance defined as *p* ≤ 0.05 and gene sets containing 2–200 genes. Normalized enrichment scores (NES) were plotted across time points.

### Literature comparison

Gene identifiers and LogFCs were obtained from published datasets. *D. discoideum* genes were mapped to human orthologues using OMA, and mouse genes using orthogene (30, 41–44). Shared LogFCs were hierarchically clustered using Euclidean distance and complete linkage. Pairwise Spearman correlations were calculated, FDR-adjusted (≤0.05), and visualized as heatmaps. For direct comparison with Agudelo et al., genes with concordant LogFC direction across 24–48 hpi and across the UT127/UT205 conditions were identified and subjected to KEGG over-representation analysis using clusterProfiler (*p* ≤ 0.05).

### Western blotting

Proteins were separated by SDS-PAGE, transferred to nitrocellulose membranes, and immunodetected as previously described (56), using ECL Prime Blocking Reagent instead of nonfat dry milk. Signals were detected using ECL Plus and a Fusion Fx imaging system (Vilber Lourmat), and band intensities were quantified with ImageJ.

### LLOMe experiments

For membrane-repair assays, 5 × 10⁵ cells were seeded in 96-well IBIDI plates. Images were acquired using an ImageXpress Micro XL HC microscope with a 60× water-immersion objective every 5 min for 1.5 h, using three wells per condition, four fields per well, and three z-sections at 1-µm intervals. After baseline imaging, LLOMe was added at the indicated concentration and GFP-Vps32 recruitment was monitored. Three independent biological replicates were performed. For cell-death assays, wt and *atg9*-KO cells were incubated with propidium iodide (1 µg/mL) and treated with LLOMe (4.5 or 9 mM) or left untreated. PI uptake was monitored by automated imaging and quantified as the percentage of PI-positive cells.

### Image and statistical analysis

High-resolution images were acquired using a Leica Stellaris confocal microscope and processed with default Lightning deconvolution settings. Bacteria were segmented from mCherry or GFP fluorescence, and host cells from the appropriate reporter channel. Reporter recruitment was quantified by identifying fluorescence-positive structures above cytosolic background and measuring their overlap with bacteria masks. Sample sizes and *p*-values are indicated in the figure legends. Statistical analyses were performed using GraphPad Prism 10 or custom R pipelines.

## Data and materials availability

RNA-seq raw data are available through the SRA/GEO BioProject PRJNA1511298, and analysis code and processed data are available via Zenodo (https://doi.org/10.5281/zenodo.21901372) and the UNIGE bioinformatics platform. Vectors generated in this study are available from the corresponding authors upon request. All data and code required to reproduce the results are provided in the article and/or Supplementary Materials.

## Acknowledgments

We thank Dr. L. Eichinger for sharing the *atg9*-KO cells and the GFP-Atg9 expression constructs. We thank F. Leuba and L. Jradi for technical support in generating expression cell lines, the Photonic Bioimaging Center, FACS Core Facility, and iGE3 Genomics Platform at the University of Geneva, and N. Chiaruttini for assistance with image analysis. We also acknowledge the ACCESS Geneva Imaging Facility for support with high-content microscopy experiments and analysis.

---

**Fig. S1.**
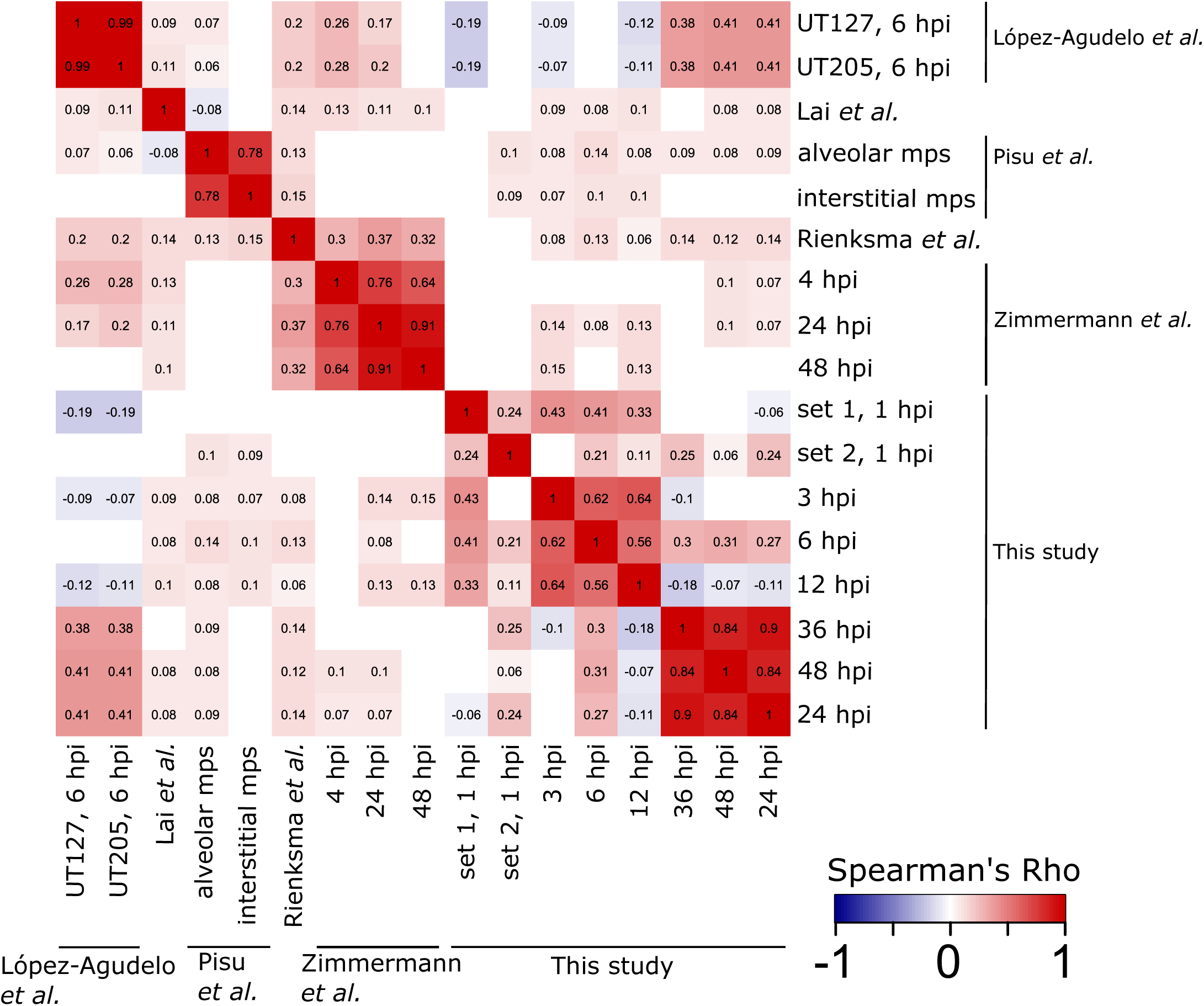
Cross-correlation analysis of various transcriptomic studies. Spearman correlation coefficients filtered by Benjamini-Hochberg corrected *p*-value ≤ 0.05 for all combinations of the selected studies. Red codes for positive correlation coefficients and thus correlation, blue codes for negative correlation coefficients and thus anti-correlation. Macrophages are abbreviated with Mps. Similarity within studies is highlighted by red blocks along the diagonal, similarity between data from this study and López-Agudelo et al. is highlighted in the upper right corner.

**Fig. S2.**
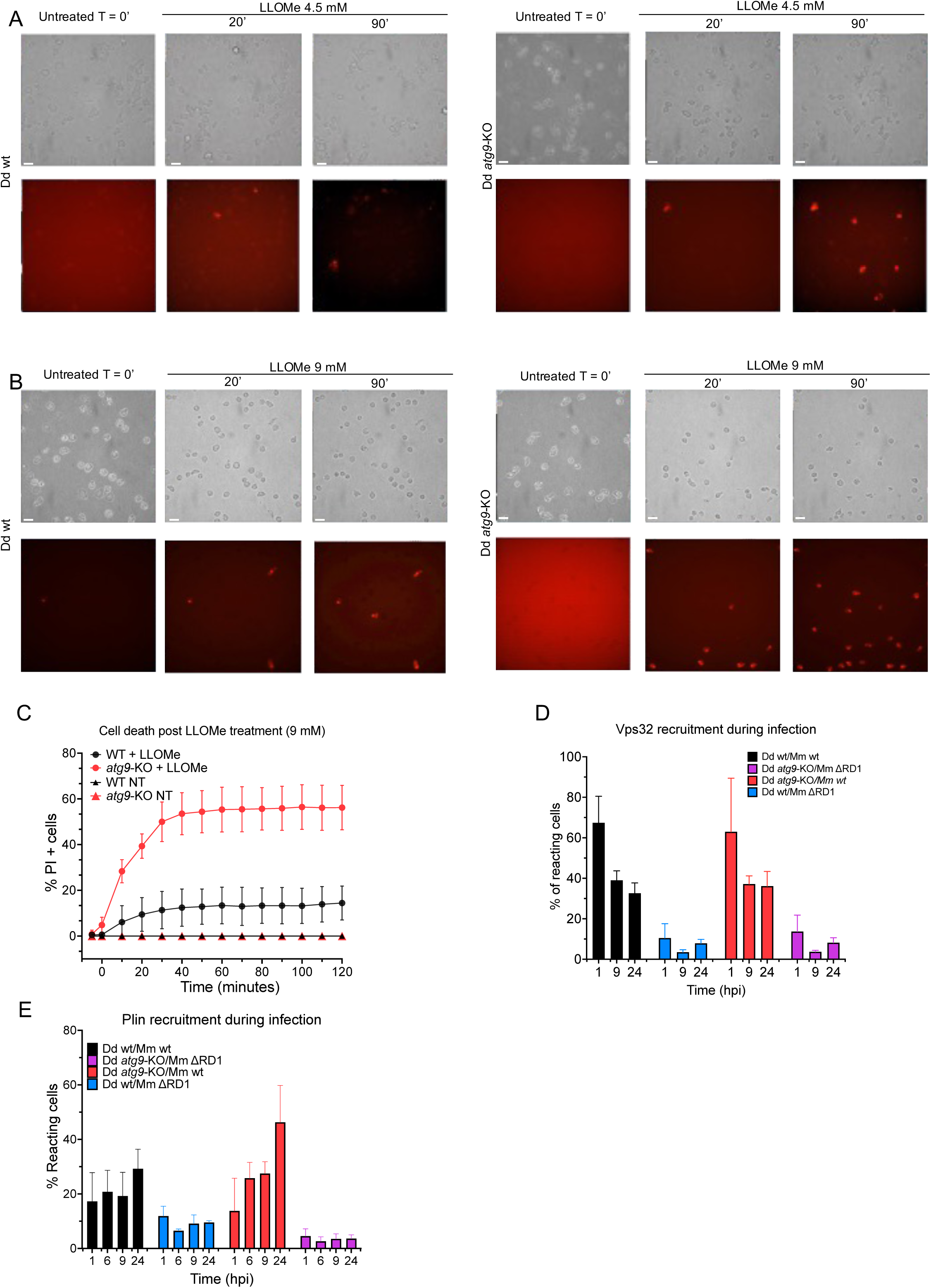
Dose-dependent membrane damage-induced cell death and quantification of GFP-Vps32 and mCherry-Plin recruitment during Mm infection. **(A)** Representative brightfield (top) and propidium iodide (PI, bottom) images of Dd wt (left) and *atg9*-KO (right) cells, untreated (T = 0’) or treated with 4.5 mM LLOMe for the indicated times (20’, 90’). PI uptake marks cells with compromised plasma membrane integrity; scale bars, 10 μm. **(B)** Representative brightfield (top) and PI (bottom) images of Dd wt (left) and *atg9*-KO (right) cells, untreated (T = 0’) or treated with a higher dose of LLOMe (9 mM) for the indicated times (20’, 90’); scale bars, 10 μm. **(C)** Quantification of PI-positive cells over time following treatment with 9 mM LLOMe in wt and *atg9*-KO cells, alongside untreated (NT) controls for each genotype. Data are presented as mean ± SEM. **(D)** Quantification of the percentage of cells positive for GFP-Vps32 recruitment at MCVs during infection with Mm wt or ΔRD1 mutant in Dd wt and *atg9*-KO host cells, at 1, 9, and 24 hpi. Data are presented as mean ± SEM. **(E)** Quantification of the percentage of cells positive for mCherry-Plin recruitment during infection with Mm wt or the ΔRD1 mutant in wt and *atg9*-KO host cells, at 1, 6, 9, and 24 hpi. Data are presented as mean ± SEM.

**Table S1: DEGs early timepoints.**

This table exceeds the size limit for upload in the submission portal, and has therefore been uploaded on Zenodo (https://doi.org/10.5281/zenodo.21901372).

**Table S2: DEGs late timepoints.**

This table exceeds the size limit for upload in the submission portal, and has therefore been uploaded on Zenodo (https://doi.org/10.5281/zenodo.21901372).

**Table S3. Literature comparison**

